# Effect of Pre-harvest Calcium Chloride and Salicylic Acid Spray on Morphological and Biochemical Traits of Guava (*Psidium guajava*)

**DOI:** 10.1101/2021.03.04.433865

**Authors:** Lochan Kaushik, Rahul Kumar, Dilip Kumar Reddy, Prashant Kaushik

## Abstract

Guava is a small, tropical fruit tree grown in various tropical and subtropical regions. Salicylic acid (SA) is a phenolic compound that enhances disease resistance and delays the fruit ripening process. Calcium is an essential cell component that delays ripening, particularly softening of the fruit. The effect of foliar spray of CaCl_2_, and SA, on the physical and biochemical traits of guava was investigated in the present investigation. The application of CaCl_2_ 2% + SA 2mM was more effective as compared with both when applied alone. The data were recorded on fruit set (%), fruit weight (g), fruit length (cm), fruit diameter (cm), fruit yield (kg), ripening period (days), TSS, acidity, total sugar, ascorbic acid, nitrogen, phosphorus, potassium. CaCl_2_ 2% + SA 2mM was showed better performance in all cases, followed by SA 2mM and CaCl_2_ 2%. Overall, this work determines the influence on guava's essential traits by pre-harvest calcium chloride and salicylic acid.

## Introduction

Guava belonging to the myrtle family (Myrtaceae) is a common fruit cultivated in tropical and subtropical regions of the world as popular fruit. The guava is full of nutrients as iron, calcium, phosphorus and vitamins as ascorbic acid, pantothenic acid, vitamin A along niacin (Embaby and Hassan, 2015). The plant's durable qualities can withstand unfavourable climatic factors and increases under a broad selection of soil sorts (Dhaliwal and Singla 2002). It is ordinarily ingested new as a dessert fruit, or perhaps refined into juice, jam, jelly, nectar, and syrup (Jagtiani et al., 1988). Salicylic acid (SA), an all-natural plant hormone, increases tolerance against biotic stresses (Khan et al., 2012). It plays a good role in plant growth, ion uptake, and nutrient transport within the plant. SA is a phenolic compound (Shafiee *et al.,* 2010), enhances disease resistance (Aghdam *et al.,* 2009), and delays fruit ripening (Srivastava and Dwivedi, 2000). SA governs processes, such as heat generation in plants, disease resistance, seed germination, gender polarisation, and ethylene production (Srivastava and Dwivedi, 2000; Ding and Wang, 2003; Zhang *et al.,* 2003). The amount of SA synthesis increased, or stress tolerance mechanisms triggered by increasing the SA concentration in the plant by external treatments have been calculated in stress-tolerant plants. (*Ding et al.,* 2001; Hayat *et al.,* 2010).

Salicylic acid increases shoot length, the number of leaves and leaf area in guava (Jamali *et al*. 2011; Ram *et al*., 2016). Foliar spray of SA significantly increased vegetative growth, several flower clusters in strawberry cultivars (Lo’ay and El Khateeb, 2011; Mohamed *et al*., 2017). Salicylic acid treatment increases total phenol content in fruit (Giménez *et al.,* 2014; Erogul and Özsoydan, 2020).

As a portion of the cellular system, calcium plays a crucial function in preserving the cell wall (Fry 2004; Hepler and Winship 2010). Ca delays the process of maturation as well as increases longevity (Shehata *et a*l., 2009). 1 % Ca (NO_3_)_2_ treated fruit has decreased spoilage, ascorbic acid and TSS by two times under context conditions appropriately (Goutam et al., 2010). CaCl_2_ modulates improvements in fresh guava and increases guava consistency during storage (Chawla *et al*., 2010). The postharvest application of CaCl_2_ (2%) improves guava fruits' shelf life with less weight loss and high quality (Mahajan *et al.,* 2011).

A combination treatment of CaCl_2_ and SA improves plant defense systems. It enhances the storage period of apples (Zhao and Wang, 2015), increase fruit firmness, and reduce percentages of weight losses and fruit decay during storage (Kazemi *et al.,* 2011; Mostafa and Sultan, 2018). Keeping this in view, the investigation was carried out to study the effect of CaCl_2_, SA alone and in combination on physical and biochemical parameters of guava.

## Materials and methods

The present study was carried out at Agriculture Research Farm, Haryana, India, coordinates at 29.94°N 76.89°E during 2018-19. The experiment was conducted on 5-year-old guava trees of variety Hisar Safeda, the plants were procured from the CCS Haryana Agricultural University, Hisar, India. The experiment was carried out in an entirely randomized block design composed of 3 replicates and the soil characters are defined, elsewhere (Kaushik, 2020). The investigation was carried out with four treatments with three replications. The treatments were comprised of Calcium chloride (CaCl_2_) @ 2%, Salicylic acid (SA) 2mM and CaCl_2_2% + SA 2mM along with control. At the colour stage (20 days before harvest) in the 2nd week of December, treatments were applied in liquid spray until the leaves were completely soaked. Each treatment consisted of 5 plants in a single replication, and there were 3 replications of each treatment.

The data were recorded on fruit set (%), fruit weight (g), fruit length (cm), fruit diameter (cm), fruit yield (kg), ripening period (days), TSS, acidity, total sugar, ascorbic acid, nitrogen, phosphorus, potassium. With the assistance of an Erma Hand Refractometer (Napoca, Japan), the total soluble solids (TSS) content was measured and expressed at 20 °C temperature. Ascorbic acid, total sugar and titratable acidity were estimated by the method described by AOAC (1990).

## Data analysis

Means of each replication was subjected for the estimation of ANOVA. Duncan’s multiple range test (DMRT) was used as the posthoc analysis to estimate the mean differences. Pearsoǹs linear correlations were calculated for the determination of correlation among the traits. All numerical analysis was performed using JASP (version 0.14.1) software program.

## Results

Maximum fruit set (84.22) was observed in treatment CaCl_2_ 2% + SA 2mM followed by the CaCl_2_ 2% (72.33) and SA 2mM (79.67) as compare by the control 76.11 (Table 1). In case of fruit weight, highest was observed in the treatment CaCl_2_ 2% + SA 2mM (67.68g) followed by the CaCl_2_ 2% (60.66g) and SA 2mM (60.02g) as compare to the control (44.78) (Table 1). The treatment CaCl_2_ 2% + SA 2mM was more effective on the fruit weight. Maximum fruit length (5.92cm) was observed in the case of treatment CaCl_2_ 2% + SA 2mM followed by SA 2mM (5.73) and CaCl_2_ 2% (5.41) as compare to the control (4.83 cm) (Table 1). Fruit diameter (5.68cm) was maximum in the treatment of CaCl_2_ 2% + SA 2mM followed by the SA 2mM (5.53) and CaCl_2_ 2% (5.28) as compare to the control 4.96 cm (Table 1). Highest fruit yield (77 kg) was recorded in case of CaCl_2_ 2% + SA 2mM spray followed by SA 2mM (62.78kg) and CaCl_2_ 2% (57.56kg) as compare to control (53.11kg) (Table 1).

**Table 1:** Effect of CaCl_2_, SA alone and in combination on yield parameters of guava.

| Treatments | Fruit set | Fruit weight (g) | Fruit length (cm) | Fruit diameter (cm) | Fruit yield (kg) |
| --- | --- | --- | --- | --- | --- |
| CaCl<sub>2</sub> 2% | 72.33c | 60.66b | 5.41 | 5.28b | 57.56b |
| SA 2mM | 79.67b | 60.02b | 5.73 | 5.53ab | 62.78ab |
| CaCl<sub>2</sub> 2% + SA 2mM | 84.22a | 67.68a | 5.92a | 5.68a | 77.00a |
| Control | 76.11bc | 44.78c | 4.83c | 4.96c | 53.11b |
| CV | 2.64 | 3.40 | 3.20 | 2.43 | 11.98 |

\*values in a column followed by the same letter are not significantly different, $p \leq 0.05$ , LSD.

Minimum ripening day (11.44) was observed in case of CaCl_2_ 2% + SA 2mM spray followed by SA 2mM (12.44) and CaCl_2_ 2% (18.56) as compare to the control (22.22) (Table 2). Highest TSS (11.12) was observed in case of CaCl_2_ 2% + SA 2mM spray followed by SA 2mM (10.87) and CaCl2 2% (10.16) as compare to the control (8.94) (Table 2). Minimum acidity (0.44) was recorded in case of CaCl_2_ 2% + SA 2mM spray followed by SA 2mM (0.47) and CaCl_2_ 2% (0.49) as compare to the control (0.51) (Table 2). Highest sugar recorded in case of CaCl_2_ 2% + SA 2mM (7.20) spray followed by SA 2mM (6.86) and CaCl_2_ 2% (6.24) as compare to the control (5.50). Highest ascorbic acid (8.29) recorded in case of CaCl_2_ 2% + SA 2mM spray followed by SA 2mM (8.01) and CaCl_2_ 2% (7.54) as compare to the control (7.24) (Table 2).

**Table 2:** Effect of CaCl_2_, SA alone and in combination on bio-chemical characters of guava.

| Treatments | Ripening day | TSS (°Brix) | Acidity (%) | Total sugar | Ascorbic acid (mg/100 g Pulp) |
| --- | --- | --- | --- | --- | --- |
| CaCl<sub>2</sub> 2% | 18.56b* | 10.16b | 0.49a | 6.24b | 7.54c |
| SA 2mM | 12.44c | 10.87a | 0.47b | 6.86a | 8.01b |
| CaCl<sub>2</sub> 2% + SA 2mM | 11.44c | 11.12a | 0.44c | 7.20a | 8.29a |
| Control | 22.22a | 8.94c | 0.51a | 5.50c | 7.24d |
| CV | 3.64 | 3.11 | 2.06 | 4.22 | 1.19 |

\*values in a column followed by the same letter are not significantly different, $p \leq 0.05$ , LSD.

## Correlation

Correlation studies were revealed that fruit set content was positively correlated with the total sugar, ascorbic acid, nitrogen and phosphorous (Figure 1). Fruit weight was positively correlated with the fruit length, fruit diameter, TSS, total sugar, ascorbic acid, and potassium (Figure 1). In comparison, a significant and positive correlation was determined between the fruit weight, fruit diameter, TSS, total sugar, ascorbic acid, and potassium with that of fruit length. Fruit diameter was positively correlated with the fruit weight, fruit length, TSS, total sugar, ascorbic acid, and potassium (Figure 1).

**Figure 1.**
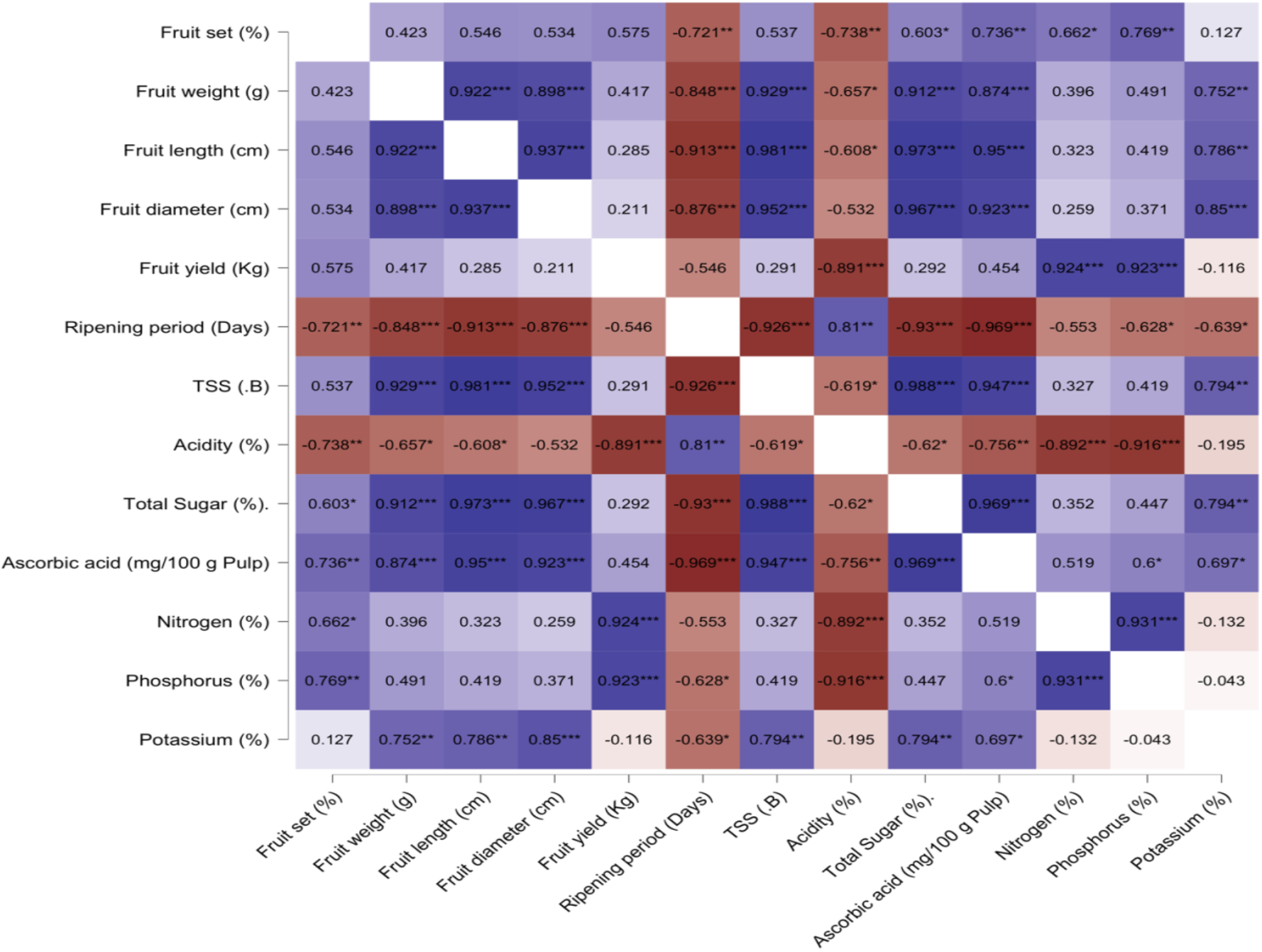
Pearson’s linear correlation among the traits studied in guava under the influence of CaCl_2_ and SA.

Moreover, the ripening period was significant and negatively correlated with all parameters except acidity. Furthermore, TSS was determined to be positively correlated with the fruit weight, fruit length, fruit diameter, total sugar, ascorbic acid, and potassium whereas, the acidity was correlated with the ripening period (Figure 1). Moreover, total sugar was positively correlated with fruit set, fruit weight, fruit length, fruit diameter. The positive correlation was determined between ascorbic acid and fruit set, fruit weight, fruit length, fruit diameter, TSS, total sugar, nitrogen, phosphorous (Figure 1). Nitrogen was significant and positively correlated with fruit set, fruit yield, whereas phosphorous was correlated with fruit set, fruit yield, nitrogen, ascorbic acid. Moreover, potassium was positively correlated with fruit weight, fruit length, fruit diameter, TSS, total sugar, ascorbic acid (Figure 1).

The highest percentage of leaf N content was recorded for the treatment comprising of CaCl_2_ 2% + SA 2mM, followed by the treatment with SA 2mM and CaCl_2_ 2% (Table 3). Similarly, the highest percentage of P content was determined for the treatment CaCl_2_ 2% + SA 2mM, followed by the treatments with SA 2mM and CaCl_2_ 2% (Table 3). Whereas the K content was highest in the treatment SA 2mM, the least values were recorded in control (Table 3).

**Table 3:** Effect of CaCl_2_, SA alone and in combination on nutrients of leaf in guava.

| Treatments | Nitrogen (%) | Phosphorous (%) | Potassium (%) |
| --- | --- | --- | --- |
| CaCl<sub>2</sub> 2% | 1.76b | 0.12c | 0.75a |
| SA 2mM | 1.79b | 0.13b | 0.79a |
| CaCl<sub>2</sub> 2% + SA 2mM | 1.88a | 0.15a | 0.78a |
| Control | 1.75b | 0.12c | 0.63b |
| CV% | 1.37 | 2.64 | 6.12 |

\*values in a column followed by the same letter are not significantly different, $p \leq 0.05$ , LSD.

Fruit set was observed maximum in the treatment CaCl_2_ 2% + SA 2mM application followed by SA 2mM CaCl_2_ 2%. Similarly, Ram *et al.,* (2016) reported that maximum fruit set per cent was found in SA @ 100 ppm. Similar result of SA on fruit set was also recorded by Nicholas and Embree (2004) and Liao *et al.* (2006) in apple and citrus, respectively. Fruit weight, fruit length and fruit diameter were highest recorded in the combination of CaCl_2_ 2% + SA 2mM. Erogul and Özsoydan (2020) also reported that fruit weight, fruit width and fruit length were increased with the application of SA. The combined effect of CaCl_2_ 2% + SA 2mM gave the highest fruit yield, followed by SA 2 mM. Our observations were similar with the findings of Kazemi (2013). Highest TSS, ascorbic acid was observed in the case of CaCl_2_ 2% + SA 2mM spray followed by SA 2mM and CaCl_2_ 2% compared to the control. Zhao and Wang (2015) also reported the similar trend on total soluble solids content, titratable acidity content. Mirdehghan and Ghotbi (2014) also reported SA and CaCl_2_ application increases TSS. Qureshi *et al.,* (2013) also described that SA+CaCl_2_ treatment improved strawberry quality by significantly increasing ascorbic acid and total soluble solids contents, with a non-significant result for titratable acidity. The data revealed that the application of CaCl_2_ 2% + SA 2mM significantly increased nitrogen, phosphorous, and potassium content compared to control. The increase of nitrogen and phosphorous were non-significant with the application of calcium chloride as compared to control (Kaushik *et al*., 2020b, 2020a). The results are also agreeing with the findings of Youssef *et al.,* (2017).

## Conclusion

The application of CaCl_2_ 2% + SA 2mM was more effective and showed more fruit yield and fruit set followed by SA 2mM and CaCl_2_ 2%. High TSS, total sugar and ascorbic acid was found in the combined effect of CaCl_2_ 2% + SA 2mM followed by SA 2mM and CaCl_2_ 2%. These observations revealed that the combined effect of SA 2mM and CaCl_2_ 2% was more effective.

## Notes

### Competing Interest Statement

The authors have declared no competing interest.

